# VizCNV: An integrated platform for concurrent phased BAF and CNV analysis with trio genome sequencing data

**DOI:** 10.1101/2024.10.27.620363

**Authors:** Haowei Du, Ming Yin Lun, Lidiia Gagarina, Michele G Mehaffey, James Paul Hwang, Shalini N. Jhangiani, Sravya V. Bhamidipati, Donna M. Muzny, M Cecilia Poli, Sebastian Ochoa, Ivan K. Chinn, Anna Linstrand, Jennifer E. Posey, Richard A Gibbs, James R. Lupski, Claudia M. B. Carvalho

## Abstract

**Background:** Copy number variation (CNV) is a class of genomic Structural Variation (SV) that underlie genomic disorders and can have profound implications for health. Short-read genome sequencing (sr-GS) enables CNV calling for genomic intervals of variable size and across multiple phenotypes. However, unresolved challenges include an overwhelming number of false-positive calls due to systematic biases from non-uniform read coverage and collapsed calls resulting from the abundance of paralogous segments and repetitive elements in the human genome.

**Methods:** To address these interpretative challenges, we developed VizCNV. The VizCNV computational tool for inspecting CNV calls uses various data signal sources from sr-GS data, including read depth, phased B-allele frequency, as well as benchmarking signals from other SV calling methods. The interactive features and view modes are adept for analyzing both chromosomal abnormalities [e.g., aneuploidy, segmental aneusomy, and chromosome translocations], gene exonic CNV and non-coding gene regulatory regions. In addition, VizCNV includes a built-in filter schema for trio genomes, prioritizing the detection of impactful germline CNVs, such as *de novo* CNVs. Upon computational optimization by fine-tuning parameters to maximize sensitivity and specificity, VizCNV demonstrated approximately 83.8% recall and 77.2% precision on the 1000 Genome Project data with an average coverage read depth of 30x.

**Results:** We applied VizCNV to 39 families with primary immunodeficiency disease without a molecular diagnosis. With implemented build-in filter, we identified two *de novo* CNVs and 90 inherited CNVs >10 kb per trio. Genotype-phenotype analyses revealed that a compound heterozygous combination of a paternal 12.8 kb deletion of exon 5 and a maternal missense variant allele of *DOCK8* are likely the molecular cause of one proband.

**Conclusions:** VizCNV provides a robust platform for genome-wide relevant CNV discovery and visualization of such CNV using sr-GS data.

## Introduction

Copy number variation (CNV), a class of structural variation (SV) that results in deviations from the diploid state, represents a major source of genetic diversity. Such CNVs have significant implication for human health and contribute to a variety of genomic disorders [1]. CNVs contribute to the allelic architecture of genes for a multitude of rare disease traits and common disorders [2]. These chromosomal and gene structural genomic alterations range from whole chromosome duplications, i.e. trisomies, to changes affecting a single exon, with pathogenic CNVs changing the expression of dosage-sensitive genes at loci for autosomal dominant (AD) traits or contributing to biallelic variation at loci for autosomal recessive (AR) traits.

The detection of CNVs has evolved significantly, from karyotyping large CNVs to more high-throughput methods to detect dosage like array comparative genomic hybridization (aCGH) and genome-wide single nucleotide polymorphism (SNP) arrays. Modern next-generation massively parallel DNA sequencing, including exome sequencing (ES) and genome sequencing [GS; both short read (sr) and long read (lr)-GS], may provide even finer scale resolution. However, CNV calling is challenged by the multitude of false positive calls particularly for copy number gains and sizes less than 50 kb. This problem may be amplified by the overwhelming number of SV calls from GS [3]. The integration of B-allele frequency (BAF) information and haplotype phasing to the CNV analysis has been pivotal in high-impact variant prioritization [4], and enabled the identification of: i) *de novo* CNVs – i.e. mutations present only in the proband’s genome, but not in either parent [4], ii) homozygous deletions [5] and duplications [6], iii) variants rendered homozygous by identity by descent [7], and the chromosome phenomenon of uniparental disomy [8,9]. Such integrative approaches have significantly narrowed the search space for potential pathogenic variants, revealing novel candidate genes in rare disease (i.e. a gene in a newly proposed gene-disease or variant-trait relationship), in addition to revealing novel disease mechanisms [10,11].

Methods for concurrent whole genome haplotyping and copy number profiling through an array-based platform have shown great utility in prenatal testing and molecular diagnosis of genomic disorders [12,13], but the resolution and sensitivity of haplotyping with SNP array is region-specific, depending on the design of the probes and density of SNPs. Mother-father-proband trio GS has the potential to enhance the resolution of CNV haplotyping by enabling the calling of both common and rare SNVs within a family. The complexity of applying this analysis to GS data, due to the numerous bioinformatic steps involved, hinders its widespread adoption.

To address these challenges, we introduce VizCNV, a tool specifically designed for the concurrent BAF, CNV and SV analysis on GS data in the context of rare genetic disease research. VizCNV enhances the application of sequencing data by incorporating BAF, SV information and parental origin to streamline the process of identifying and prioritizing high-impact allele-specific CNVs. VizCNV is built on the R Shiny framework and is designed to be interactive and user-friendly.

## Methods

### Read depth normalization and segmentation

A BED file that stores the genomic position and sequencing coverage measured from BAM or CRAM file is required. While multiple tools can generate such a file, VizCNV seamlessly supported the bed file output from Mosdepth [14]. Two choices are available for normalization and segmentation in the application. By default, the chromosomal median of read depth is used to calculate the log_2_ ratio for every 1000 bp. Additionally, there is an option to use the genome-wide median of read depth, which facilitates chromosomal analysis, such as detecting aneuploidy. The log_2_ ratio data points can be segmented using the Circular Binary Segmentation (CBS) algorithm [15] implemented in the DNAcopy Bioconductor package or shift level model implemented in the SLMSuite R package [16]. Problematic regions of the genome (build GRCh38 and hg19), as defined by those non uniquely specified DNA stretches outside the ‘assayable portion’ of the human genome [17], were masked using UCSC RepeatMasker and segmental duplication (SD) tracks.

### Trio genome-wide SNV genotyping and phased BAF segmentation

Germline variants per sample, including SNVs and small insertions and deletions (INDELs), were called using the Genome Analysis Toolkit (GATK) HaplotypeCaller in ‘-ERC GVCF’ mode. This generated a genomic VCF (gVCF) file for each family member: the proband, mother, and father. Subsequently, joint genotyping was conducted on the combined gVCFs for the entire family, facilitating genotyping at all variant positions. Variants were excluded from phasing if they met any of the following criteria, i) absence of genotype information for any family member, ii) low coverage depth in any one family member (read depth < 8), and iii) presence of an INDEL or being a multiallelic site [18]. SNVs were phased confined to Mendel’s expectations. The variant read and total read ratio of phased B-allele were segmented using the CBS algorithm [15]. Since only informative heterozygous SNV were able to be phased and segmented, regions with runs of homozygosity (ROH) will be missed. Therefore, a mixture of phased and unphased SNV regardless of the zygosity were segmented with the same procedure to capture potential ROH regions.

### CNV Classification and Built-in Filter Module – FindCNV

Based on the benchmark performance on 1000 Genomes Project data, and internal GS data analysis, we included a built-in CNV calling module that will classify the segments with log_2_ ratio into heterozygous deletion [log_2_(0.45)-log_2_(0.55)] and duplication [log_2_(1.35)-log_2_(1.65)], normal [log_2_(0.9)-log_2_(1.1)], triplication [log_2_(1.8)-log_2_(2.2)] and multiple gain [log_2_ above 1.175]. Segments with log_2_ ratio outside the above ranges are classified as unknown for further assessment. In male cases, short reads in the pseudoautosomal region (PAR) on chromosome Y are mapped by default to the PAR reference region on chromosome X. This results in an increased log2 ratio at the PAR, which should not be interpreted as a biological duplication. These technical artifacts do not affect female cases. The read depth visualization will only work on the unique sequence region of chrY for male cases. Potential *de novo* CNVs in autosomes (chr1-chr22) will be annotated if the aberrant copy number segment is only present in proband genome by comparing genomic segments between proband and parental segments using ‘GenomicRanges::findOverlaps’ function. Homozygous CNV can be filtered in if in a genomic segment with triplication log_2_ ratio observed in the proband and duplication log_2_ ratio observed in both parental data, or absence of log_2_ ratio in proband and heterozygous deletion in both parental samples. After CNV classification, the built-in filter will report CNV with size > 10 kb and not overlap with gap track from UCSC genome browser. The SD track from the UCSC Table Browser was preprocessed and used to overlap with the CNV calls. Specifically, segmental duplication segments were merged if they were less than 2 kb apart.

### Phased B-allele Frequency Visualization

Allelic imbalance analysis was performed based on phased BAF. In trio sequencing, biallelic SNVs with > 7 read depth in all individuals were genetically phased based on Mendelian expectation and classified into five categories, phased to maternal (P1) allele, phased to paternal (P2) allele, non-phased homozygous allele, variant deviates from Mendelian expectation, non-informative allele. All classified variants were then segmented and visualized with log_2_ ratio in VizCNV. The segmentation is performed independently for phased B-allele on P1 and P2 site and potential ROH block on P1 and P2 site plus non-phased homozygous site with the CBS algorithm.[15]

### User interface of VizCNV

An interactive user interface is built to guide the user to upload the required files to run VizCNV. The input files come from a typical GS pipeline (Additional file 1: Figure S1a). The genomic analysis can be divided into three major views, genome-wide view, chromosomal view, and table view that allows for direct inspection of variable ranges (Additional file 1: Figure S1b).

### Short read genome sequencing

Genome sequencing was conducted at the Human Genome Sequencing Center (HGSC) at Baylor College of Medicine (BCM) as part of the Genomics Research to Elucidate the Genetics of Rare Diseases (GREGoR) initiative. Following quality control (QC), sequencing libraries were prepared using KAPA Hyper reagents and sequenced to produce 150 bp paired-end reads. The samples were organized into multiplexed pools to achieve an average coverage of 30x, utilizing the Illumina NovaSeq 6000 system. This approach resulted in an average sequencing yield of 108 Gb per sample, with 96% of the genome covered at a depth of 20x or greater. To ensure sample identify and integrity the Fluidigm SNPtrace™ method for rapidly genotyping 96 SNP sites was employed to verify gender prior to sequencing and to detect contamination. Using this assay sample identity was verified using the Error Rate In Sequencing (ERIS) software developed at the HGSC.

## Results

### Overview of the joint BAF and CNV analysis with VizCNV

To prioritize potential homozygous and *de novo* CNVs from trio sr-GS data, we developed an algorithm to call these variants based on joint BAF and CNV analysis of trio WGS data. There are multiple parameters affecting the sensitivity and specificity of the procedure, while the VizCNV allows for customization in all these parameters, we sought to establish a baseline that optimized the precision and recall and can be ‘plugged-in’ and implemented. The BAF (Fig. 1a) and read depth analysis (Fig. 1b) were tuned separately before the joint analysis (Fig. 1c). For read depth analysis, the per window read depth value were normalized, segmented, and classified into heterozygous deletion, heterozygous duplication, homozygous deletion, triplication, multiple gain normal or unknown based on log_2_ ratio. The performance of CNV classifications were then benchmarked with merged CNV calls from the 1,000 genomes project [19] to optimize parameters like mappability, range of the log_2_ ratio, and size of CNV. Predicted inheritance of the CNV were then classified by comparing the region between proband and parents (Fig. 1a). For the phased BAF analysis, we tuned the ROH and allelic imbalance calling on one of the 1000 Genomes trios (HG00405, HG00404 and HG00403). The inheritance was then reclassified again with the combined phased BAF data and CNV data.

**Fig. 1.**
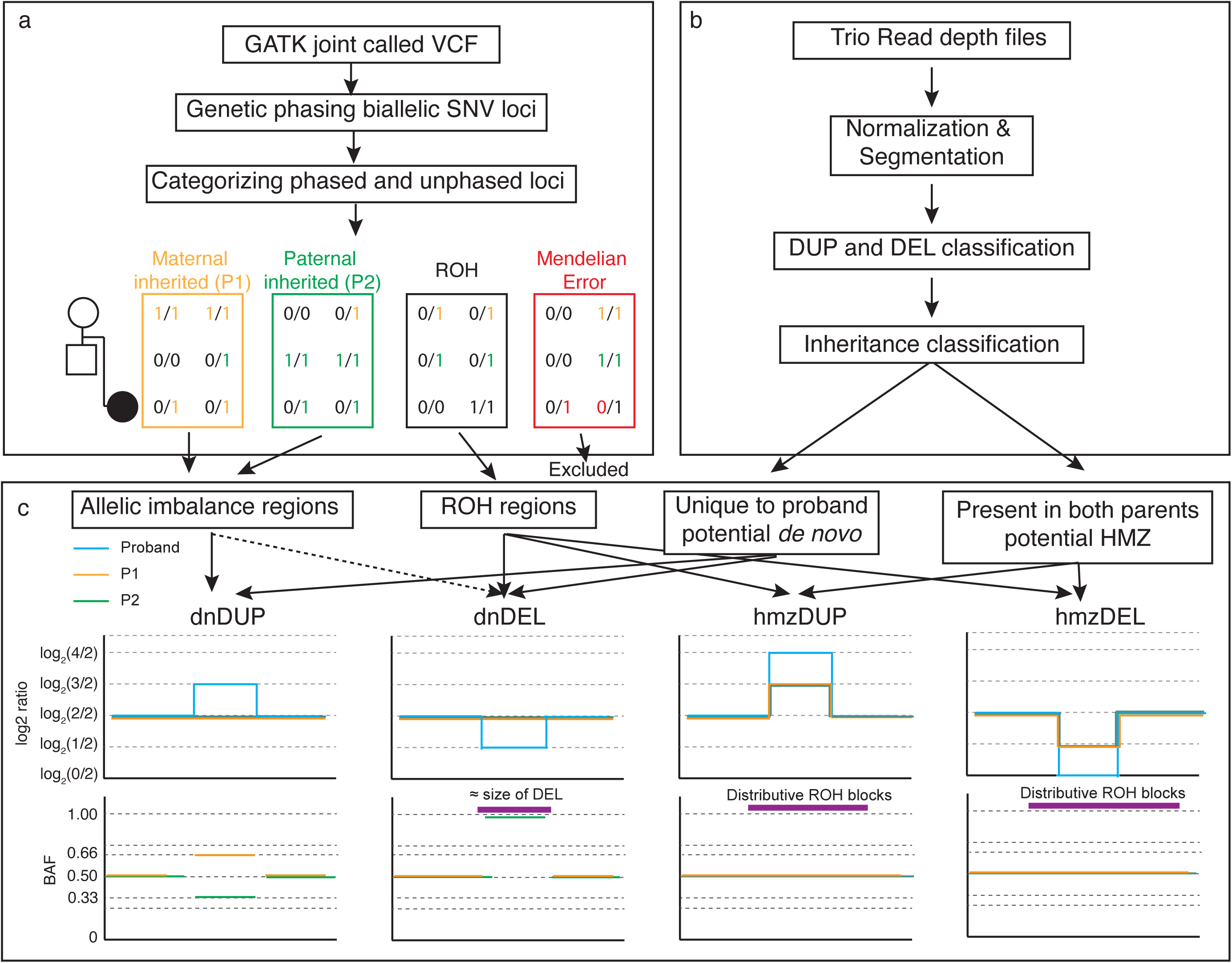
Overview of Integrating BAF and CNV Analysis Using VizCNV. **a)** Demonstrates the methodology for trio-based joint calling and the classification of B-alleles. **b)** Outlines the approach for analyzing log_2_ ratios and categorizing CNVs. **c)** Illustrates the combined analysis and visualization process using phased BAF and log_2_ ratio data for *de novo* duplication and deletion, homozygous duplication and deletion (left to right). For a *de novo* duplication occurring on the maternal allele, we would expect a log_2_ ratio of log_2_(3/2) in the proband’s read depth plot, accompanied by phased BAF values of approximately 0.66 for SNVs phased to the maternal allele and 0.33 for those phased to the paternal allele. For a de novo deletion on the maternal allele, we would anticipate a log_2_ ratio of log_2_(1/2) in the proband’s read depth plot, with the paternal phased BAF near 1, resembling a ROH similar in size to the deletion, or an allelic imbalance pattern if the deletion is mosaic. In the case of a homozygous duplication, we would expect a log_2_ ratio of log_2_(3/2) in both parents and log_2_(4/2) in the proband, with the duplication overlapping a larger ROH region. For a homozygous deletion, the expected log_2_ ratio would be log_2_(1/2) in both parents, with no signal in the proband, and the deletion would overlap a larger ROH region.

### Benchmark read depth analysis with 1000 Genome CNV calls

We selected 10 unrelated individuals from the 1000 Genomes Project (Additional file 2: Table S1) with average depth of coverage of 31.5x mapped to GRCh38, for which raw alignment files were accessible through the data portal (https://www.internationalgenome.org/data). CNV calls for these genomes were extracted from the initial study [20]. Notably, the original dataset exhibited a bias, with a significantly higher number of deletions (n = 8,584) compared to duplications (n = 54). Consequently, our benchmarking focused solely on deletion calls from these 10 sr-GS. Analysis of heterozygous deletions revealed that CNVs smaller than 7 kb demonstrated poor recall and sensitivity. Conversely, for CNVs larger than 7 kb, the highest F score was observed when applying a mappability filter of q30 (Additional file 1: Figure S2), resulting in an optimized recall of 83.8% (95% Confidence Interval (CI) [78.9%-88.7%]) and precision 77.2% (95% CI [72.3%-82.1%]).

### Phased B Allele Analysis with Trio GATK Joint Genotyping

In the joint genotyping analysis, we identified 5,019,224 biallelic single nucleotide variants (SNVs) across the autosomes (chromosomes 1-22, mapped to GRCh38) within the HG00405 family. By comparing genotypes between the proband and the parents, we classified 34% (n=1,750,252) of the SNVs as informative, attributable to one of the parents. About 54% (n= 2,725,145) of the SNVs are classified as homozygous for alternative (1/1, n=1,518,744) or reference (0/0, n=1,206,401) alleles (i.e. nonphased homozygous which indicate potential ROH signals) (Fig. 2a, b). Approximately 0.7% (n=36,123) of the SNVs deviate from Mendelian inheritance expectations. This number is significantly higher than the expected for biological *de novo* mutation rate from sr-GS [21,22] which is around 60-80 SNVs per generation. Therefore, we consider most of these deviations to be genotype errors/false discovery from the GATK joint genotype caller. (Fig. 2b). The false discovery rate (FDR) in regions that do not overlap with any segmental duplications or RepeatMasker elements is lower and estimated at 0.2%, compared to the overall rate of 0.7%. Notably, and as expected [18], regions annotated as SD or some regions annotate with RepeatMasker (e.g., SVA, Satellite, and snRNA) exhibited a significant high increase in FDR (odd ratio >10, p-value < 10^-25^) (Fig. 2c, Additional file 2: Table S2).

**Fig. 2.**
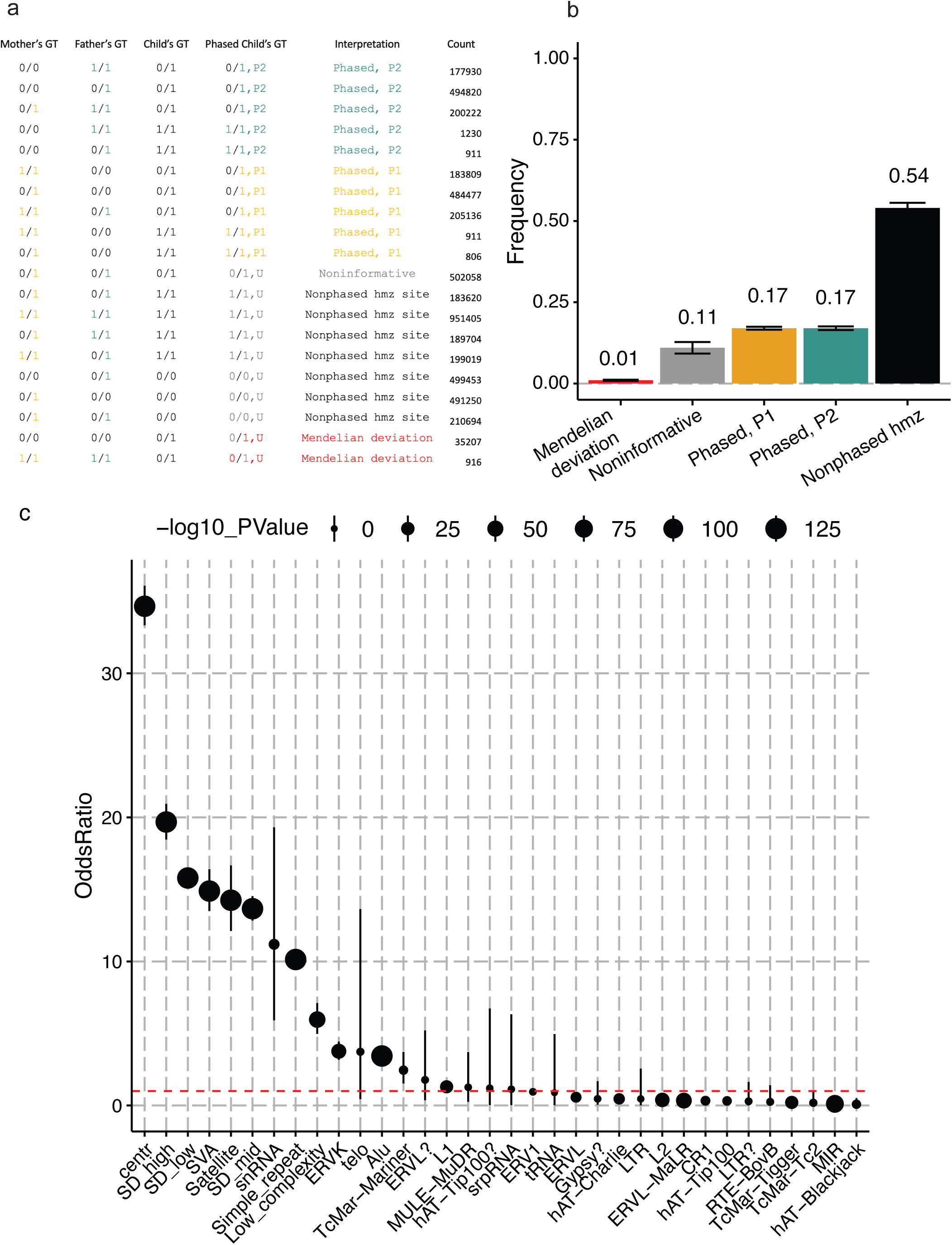
Phased BAF Analysis in a High-Coverage 1000 Genomes Project Trio. **a)** Classification principles for five categories: P1 (phased to parent 1), P2 (phased to parent 2), MV (Mendelian Violation), ROH (runs of homozygosity), and NI (Non-Informative), explaining the assignment process. **b)** The average distribution of these categories across chromosomes 1 to 22 in the trio (HG00405, child; HG00404, mother; HG00403, father), with error bars representing the 95% confidence interval. **c)** The odds ratio for the false discovery rate in regions masked for repeats and segmental duplications compared to unique genomic regions.

### Joint BAF and log_2_ Ratio Analysis on HG00405 Family

The read depth analysis of HG00405 revealed 23 heterozygous duplications and 33 heterozygous deletions with length > 10 kb in non-problematic regions (see the definition of problematic regions in the Methods section). Manual inspection of these regions identified and excluded 30 (53.6%) CNVs that map to uncharacterized problematic regions, likely representing false positive calls. Consequently, 26 CNVs were retained for further analysis. The most common false positive cases constitute CNV in regions containing two distinct segmental duplications within a 1 kb – 4 kb interval of unique sequence. Out of 26, 10 CNVs, 7 heterozygous duplications and 3 heterozygous deletions, have at least two phased SNVs from either parent, enabling joint log_2_ ratio and BAF analysis (Additional file 1: Figure S3). Of the three heterozygous deletions, the origin of the two can be inferred (one maternal, one paternal) with the average BAF of the remaining allele being 0.98 and 1 respectively. The inheritance of these two deletions were consistent with read depth analysis. Among the seven heterozygous duplications, the origin could be inferred for six (two maternal, four paternal) using read depth analysis. The one duplication that failed the read depth classification had a log_2_ ratio of 0.73 in the mother, which exceeded our set threshold for duplications [log_2_(1.35)-log_2_(1.65)], resulting in its classification as undetermined (UND) rather than as a maternal inherited duplication. The phased BAF plot independently confirmed five of the six duplications, with average BAF ranges of 0.64-0.72 on the duplicated alleles and 0.28-0.39 on the non-duplicated alleles. The exceptional duplication, approximately 1 Mb in size, had a log_2_ ratio of 0.58 in both the proband and the father, suggesting a paternally inherited duplication. The phased BAF for the duplicated alleles was 0.52 (n=705 paternal phased SNPs), while the non-duplicated alleles had a phased BAF of 0.39 (n=507 maternally phased SNPs). The segmented BAF plot shows multiple crossovers within the duplication suggesting a mixture of haplotypes in the duplicated region (Additional file 1: Figure S4).

### CNV Analysis on a pilot Primary Immune Deficiency Disease cohort [N = 39 trios]

We implemented the VizCNV pipeline in a cohort of 39 probands, who were previously ‘unsolved’ for a potentially contributing pathogenic variant with exome sequencing (ES); this cohort of subjects each had clinical evidence implicating a primary immunodeficiency disease (PIDD). Through chromosomal abnormality analysis with VizCNV, we identified a trisomy 21 case (BH10262-1) which was initially observed by clinical CMA analysis; thus, acting as positive control for the algorithm for chromosomal analysis using genome-wide median option for normalization. Detailed BAF and log_2_ ratio inspection strongly suggested a meiosis I non-disjunction event (Additional file 1: Figure S5).

VizCNV reported 6,243 inherited CNVs and 7 potential *de novo* CNVs > 10 kb in size that mapped to unique genomic intervals from the PIDD cohort (n=39 trios) (Fig. 3). Further refinement using an internal curated PIDD gene list narrowed this number to 48 rare inherited CNVs (i.e. frequency < 0.05 in gnomAD SV 2.1 database). Classification as likely true positive CNVs was based on Integrative Genomics Viewer (IGV) inspection (Fig. 3) for read depth from proband and parent samples data and breakpoint junction analysis of soft-clipped reads when they are present. Of the 48 likely true positive CNVs in 39 probands, 28 consisted of heterozygous duplications whereas 11 are heterozygous deletions. Each CNV had at least two phased SNVs from either parent, creating a reliable list for evaluating the performance of log_2_ ratio and BAF analysis (Additional file 2: Table S3). Among the 11 identified heterozygous deletions, 7 exhibited log_2_ ratio within the expected range for heterozygous deletion [log_2_(0.45)-log_2_(0.55)], classifying them as heterozygous deletion (HET_DEL). The remaining 4 deletions, with log_2_ ratio outside this range, were classified as undetermined (UND). The average BAF present in the non-deleted alleles ranged from 0.74 to 1.00. Of the 28 identified heterozygous duplications, 20 showed the presence of both paternal and maternal SNV, while 8 exhibited only one origin (either paternal or maternal). Among the 20 duplications with duplicated informative phased SNPs for both parents, 11 had BAF ranging from 0.62 to 0.70 or 0.29 to 0.37. Of these 11, 10 duplications were consistent with the inheritance inferred from read depth analysis, with one exception identified in case BH9703-1(Additional file 1: Figure S6 and Additional file 2: Table S4). This exceptional duplication had an average BAF of 0.68 (9 maternal origin SNVs) and 0.33 (21 paternal origin SNVs) (Additional file 2: Table S5). However, the log_2_ ratio analysis and IGV junction analysis suggested that this duplication was paternally inherited. Different from what was observed in the exceptional HG00405 duplication, manual inspection of the phased BAF plot did not find ratio switches within the duplication region (Additional file 1: Figure S7 and Additional file 2: Table S5).

**Fig. 3.**
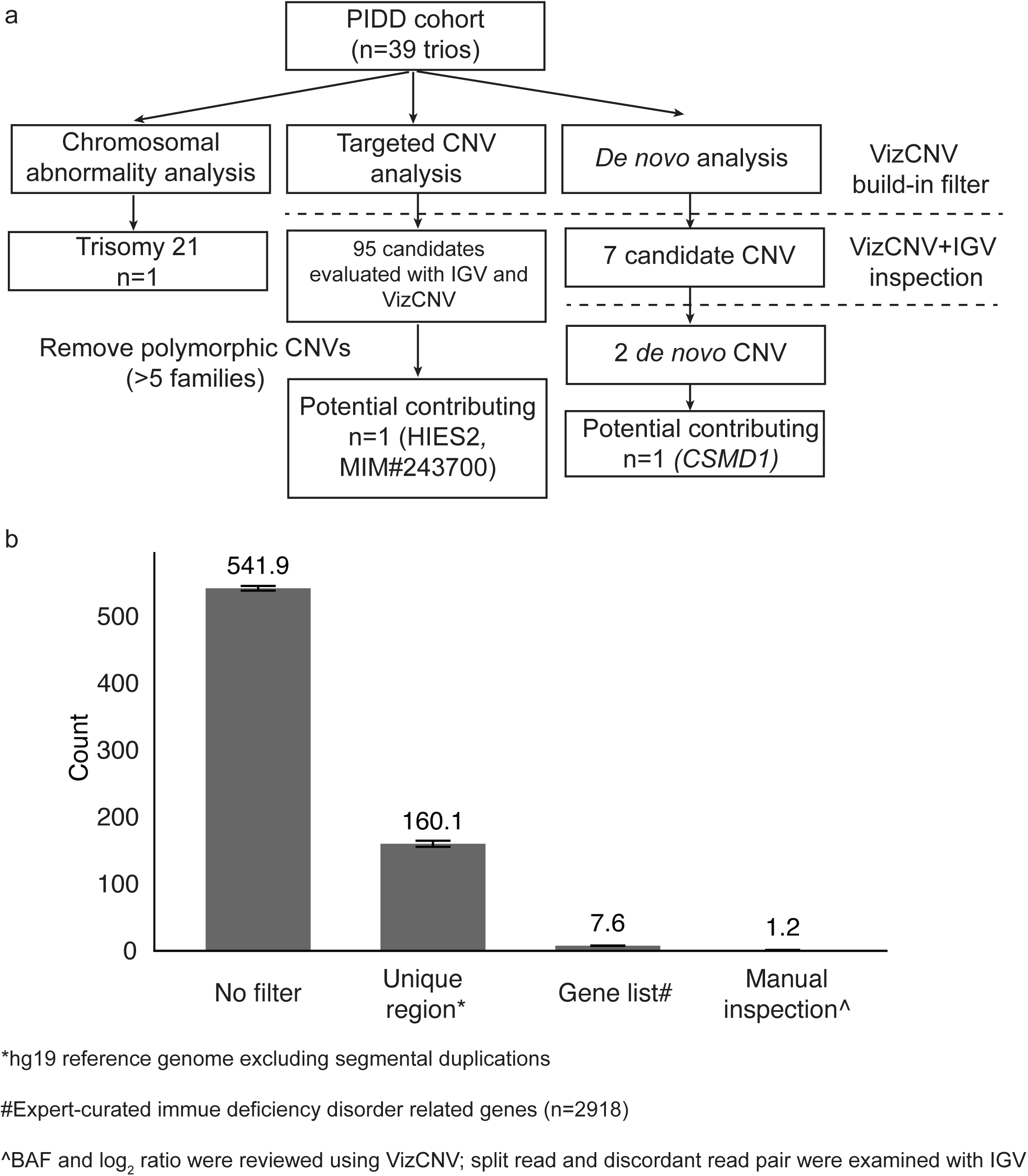
Systematic Workflow for CNV Analysis in PIDD Cohort with Stepwise Filtering and Manual Inspection. **a)** Detailed workflow for the genomic analysis in PIDD cohort consisting of 39 trios. Targeted CNV analysis identified 95 candidate CNVs, which were filtered by removing polymorphic CNVs present in more than five families, resulting in one potential contributing CNV (*DOCK8*, MIM#243700). *De novo* analysis identified seven candidate CNVs, which were further inspected using VizCNV and IGV, narrowing down to two *de novo* CNVs, with one potential contributing CNV (*CSMD1*). **b**) Average count of CNV candidates per family progressing through each stage of the selection process.

### Genotype and Phenotype Analysis

Our targeted CNV analysis that focused on curated immunodeficiency genes uncovered a paternally inherited 12.8 kb exonic deletion including exon 5 of *DOCK8* (Fig. 4). This deletion CNV is present in both proband (BH9040-1) and father (BH9040-3). The deletion was not identified in the initial ES analysis, highlighting the critical advantage of sr-GS in detecting single exon CNVs. The breakpoints of the deletion are within two *Alu* elements, with 24 bp microhomology at the breakpoint junction, supporting an *Alu*-*Alu* mediated rearrangement (AAMR) as the mutational mechanism [23]. Of note, the AluAluCNVpredictor score for *DOCK8* is 0.573; multiple studies have reported different exon deletion alleles for this gene in association with Hyper-IgE syndrome [24–27]. Rare SNV analysis on ES data identified a maternal inherited and ultrarare missense variant (NM_203447.4:c.5599T>G, p.(Phe1867Val) in *DOCK8.* The SNV is present only in this family (BH9040-1 and BH9040-2) in the Genetics of Rare Diseases (GREGoR) and absent in control databases (gnomAD v4.0, MCPS [28], and All of Us). Moreover, the amino acid position exhibits significant evolutionary conservation (PhyloP100way score 7.781). The clinical presentation of the case included elevated IgE levels, IgM deficiency, recurrent infections with herpes viruses and Staphylococcus spp, and T cell deficiency, phenotypically consistent with autosomal recessive Hyper-IgE syndrome (MIM#243700).

**Fig. 4.**
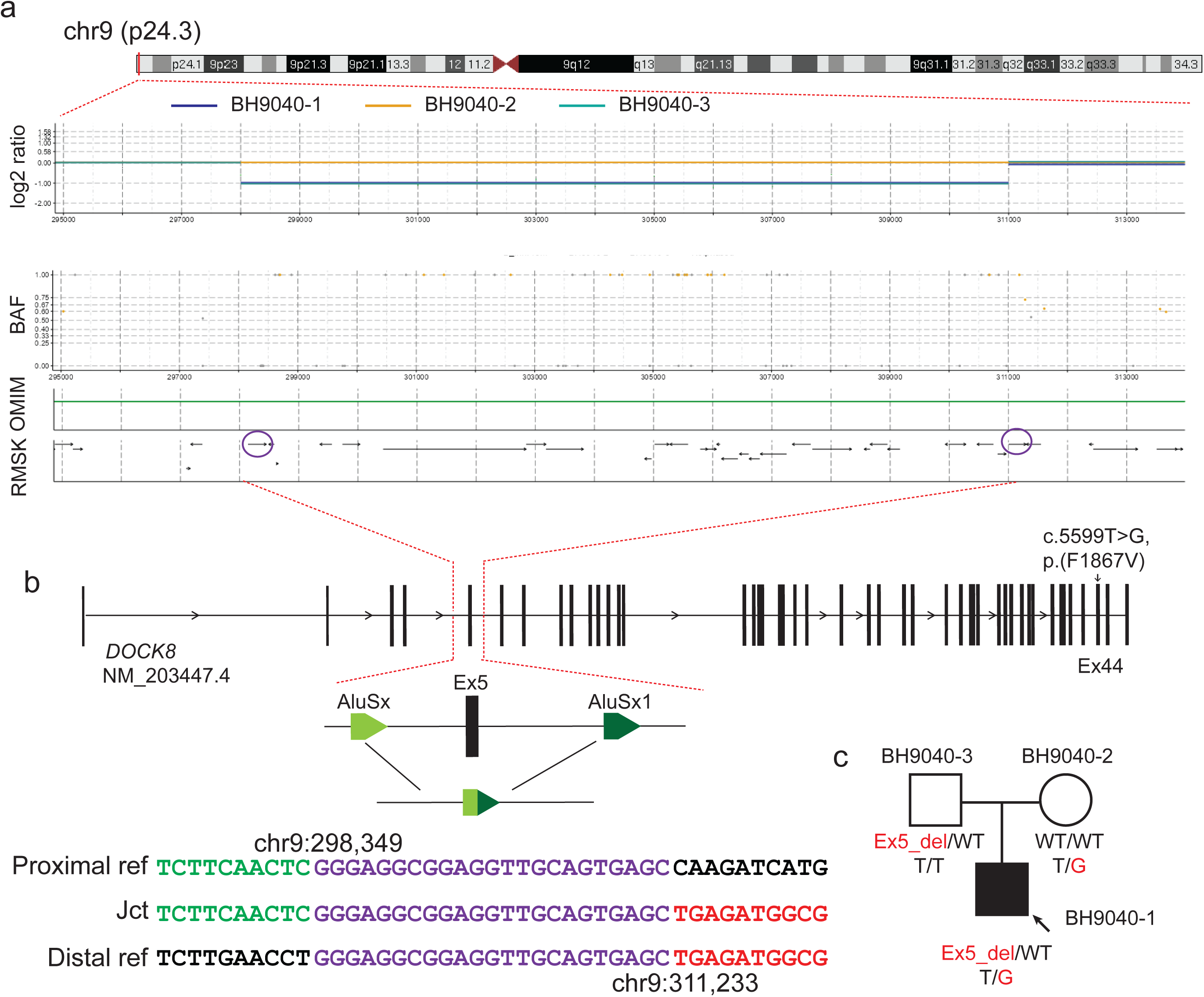
Compound heterozygous variants on *DOCK8*. **a)** On top shows the karyogram of chr9, below shows the screenshot of VizCNV chromosomal view including read depth and B-allele frequency plots showing paternally inherited a heterozygous deletion of exon 5 in *DOCK8.* The read depth graph (log_2_ ratio plot) shows the heterozygous deletion in the proband and father. Blue: proband, Yellow: mother, Turquoise: father. The BAF plot correlates with the deletion, showing a lack of heterozygous SNPs and maternally phased homozygous B alleles near 1, aligning with the paternal deletion. The OMIM track shows that this variation locates in *DOCK8* gene. The bottom track shows that repeats elements distribute across the region. The purple circles highlight the *Alu* elements overlapping with breakpoints. **b)** On the top shows the transcript structure of *DOCK8,* noting a maternally inherited missense mutation (NM_203447.4:c.5599T>G, NP_982272.2:p.F1867V) in exon 44 marked. A detailed view of the deletion’s genomic architecture is provided, alongside the breakpoint junction sequence alignment, which suggest an *Alu-Alu* mediated deletion. **c**) Segregation of the variant within the family (BH9040).

For the seven *de novo* CNV calls from 39 families, subsequent manual verification with IGV and VizCNV joint BAF and log_2_ ratio inspection confirmed three high-confidence *de novo* CNVs – two deletions and one duplication. Particularly noteworthy is the identification of two *de novo* CNVs, one deletion and one duplication, from two different families (BH8461 and BH14227) occurring within the same genomic regions (chr11:18944000-18964000, ∼20 kb in size); these are flanked by two LINE elements, hinting at a potential recurrent CNV potentially by LINE-LINE recombination. In addition to the two *de novo* CNVs, eighteen inherited CNV were identified at the same locus. The recurrent CNV predicted deleted or duplicated gene *MRGPRX1.* The other *de novo* deletion (∼59 kb) was identified within intron 3 (NM_033225.6) of the gene *CSMD1* in BH9703-1. The deletion does not remove any exons of *CSMD1* but rather removes a non-coding element ENSG00000286934 which encodes an antisense RNA to *CSMD1* (Additional file 1: Figure S8).

#### Models explain the read depth and BAF analysis difference

We propose different models to explain the observed log_2_ ratio and BAF patterns for duplications (Fig. 5). Consistent log_2_ ratio and origin of BAF for inherited alleles were observed in 5 out of 6 duplications in HG00405 family and 10 out of 11 duplications in PIDD cohort (Fig. 5a). For the ∼1 Mb duplication observed in HG00405 with inconsistent parent-proband read depth and origin of BAF alleles, we suggest a multi-step model. Initially, the duplication likely resulted from a *de novo* event from the paternal family. Subsequently, the mixture of haplotypes within this duplication arose from multiple recombination events, which disrupted the linkage of the haplotype blocks (Fig. 5c). In contrast, for the second duplication (BH9703-1), no haplotype switches were identified within the duplication region. However, one of the duplicated haplotypes was completely different from the others (Additional file 1: Figure S7a), indicating a potential interchromosomal duplication mutagenesis in contrast with an intrachromosomal duplication which seems to reflect majority of the events observed in this cohort and others [29]. This implies a single-step process where the duplicated segment might have originated from homologous chromosomes, possibly during meiosis, leading to the observed discrepancy (Fig. 5b).

**Fig. 5.**
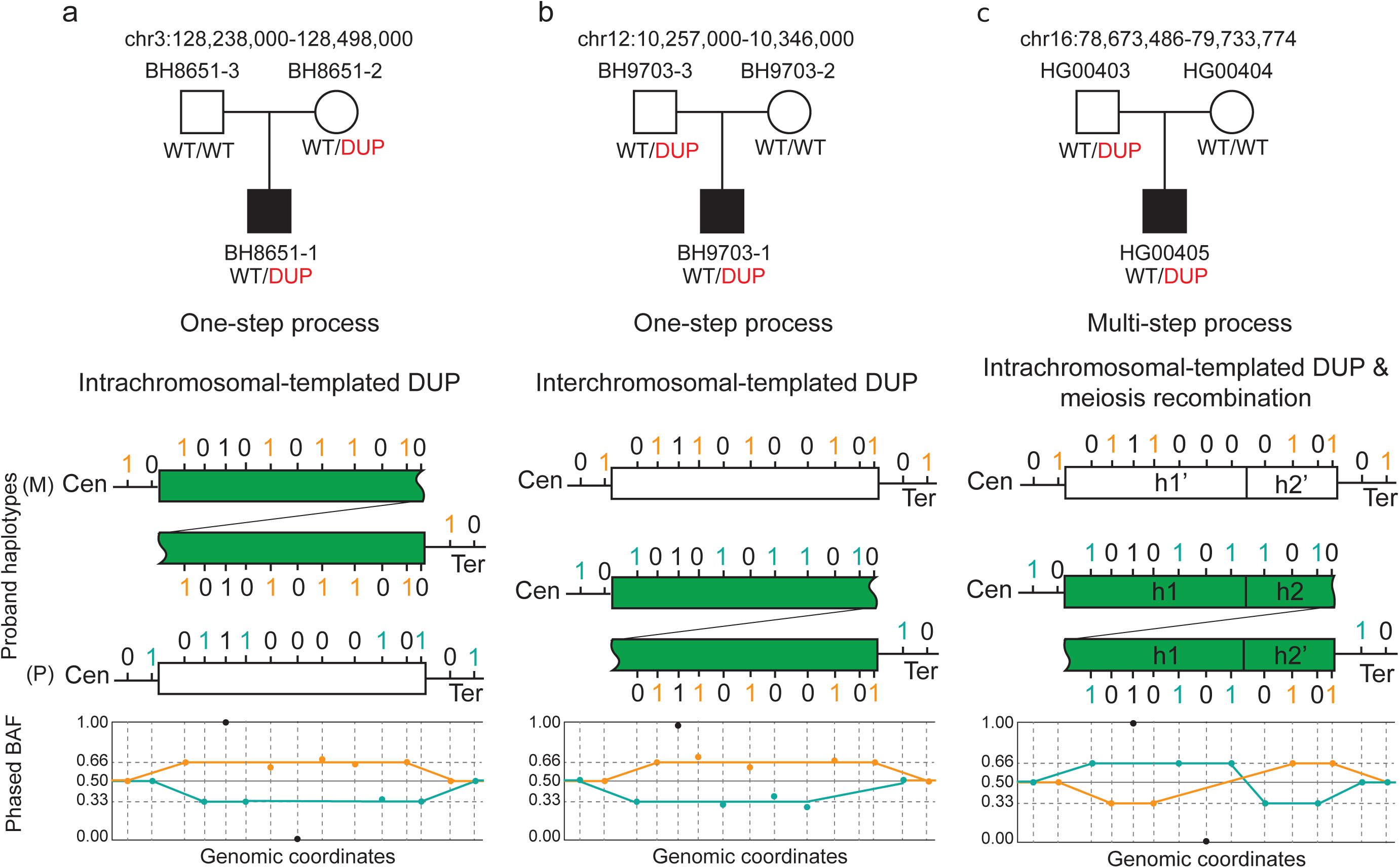
Proposed Models Explaining Inconsistencies Between Read Depth and BAF Inheritance Analysis. The top panel shows the pedigree with duplication segregation information inferred from read depth and IGV analysis. Below the pedigree are the proposed models explaining the observed BAF patterns. The two lines and rectangular segments represent two distinct haplotypes in the proband: one paternally inherited (P) and one maternally inherited (M). The green rectangle indicates the tandem duplication in the paternally inherited segment. Tick positions mark informative SNPs, with colors indicating phasing information: maternal phase (orange) and paternal phase (teal). The bottom section shows the phased B-allele frequency plot**. a)** Single-step model with intrachromosomal duplication; read depth analysis suggests maternally inherited duplication with expected phased BAF of maternal (0.66) and paternal (0.33) SNPs. **b)** Single-step model; read depth analysis suggests paternally inherited duplication but with inconsistent phased BAF of maternal (0.66) and paternal (0.33) SNPs, suggesting potential interchromosomal duplication. **c)** Multi-step model with intrachromosomal duplication and phased BAF swap, indicating a mixture of haplotypes in the duplication region due to meiosis recombination.

## Discussion

Manual inspection, often through tools like the IGV in trio analyses of proband and parental genomes, is essential to assess the structure, origin, and genomic context of CNVs. This is critical for the interpretation of the variant allele’s potential consequence to gene structure or mRNA expression [10,11], in hypothesizing SV mutagenesis mechanisms [30], and organismal hypermutation processes [4,31]. As the field of CNV analysis with sr-GS data progresses, it must continually adapt to overcome issues such as false positive variant calls and the dissection of complex genomic rearrangements (CGRs). This may challenge the determination of the parental origin – maternal versus paternal, or time in organismal development (e.g. post-zygotic) – both crucial inferences for understanding the impact of these CNV.

Tools have been recently developed to visualize SV that greatly reduced the false positive calls [32]. The application offers a range of customizable parameters that can be tailored to different types of data and research questions. These features make VizCNV a valuable and accessible tool for rare disease research, particularly for those without specialized bioinformatics expertise. The other advantage is that VizCNV combines multimodal data, which allows the visualization of the results of orthogonal techniques, which is important for validating the hypothesis and genetic model when studying rare diseases. A unique built-in feature of VizCNV enables one to visualize the chromosomal origin of the mutational event, a feature that was initially developed to help understand hypermutation events [4,31] and is essential to understanding SV mutagenesis mechanisms.

Allelic imbalance can be introduced by multiple mutational events, i.e., non-segregation of mitosis, such as intrachromosomal duplication [29,33]. The synergistic analysis of BAF and CNV greatly extends the utility of sequencing data beyond the identification of mutation. The annotation of phased SNVs with log_2_ segments facilities inferring the process of inheritance and genomic rearrangement (e.g., concurrence of recombination and CNVs, complex chromosomal rearrangement [CCR], or *de novo* CNV involving interchromosomal template switching). There is not a single model that is able to capture all these scenarios. For more complex cases, the interactive features of VizCNV can identify inconsistencies and support further mechanistic analysis to help interpreting Mendelian inconsistencies.

Fine-tuned parameters for a wide range of CNV detection could be cost prohibitive, given the space parameters (size, clonality, read depth, etc.), with target search (e.g., knowing a loss of function allele for an AR gene, looking for another potential loss of function or gain of function allele allows for a narrowed search). VizCNV provides such a resource to examine for the region and with additional statistical support.

One limitation of VizCNV is compatibility with different sequencing platforms. VizCNV has been optimized for sr-GS data, but it may not be fully compatible with lr-GS or all sequencing technologies. This could limit the applicability of VizCNV to certain rare disease research contexts. Ongoing developments include a long read specific pipeline in the future that incorporates VizCNV with epigenetic signatures. Another limitation of our study is the challenge of integrating multiallelic sites into the haplotype inference process with VizCNV. Multiallelic sites hold potential for distinguishing complex haplotype origins and segregations, which requires more sophisticated analytical tools and design [18]. Incorporating these sites could significantly enhance our understanding of complex genomic variations and their implications in genetic disorders.

### Conclusions

VizCNV has the potential to be a valuable resource for researchers and clinical genomicists worldwide. Overall, VizCNV may improve our understanding of genetic mechanisms underlying structural variation, and the clinical diagnosis and management of rare genetic diseases.

### External resources

UCSC annotation resources: https://genome.ucsc.edu/cgi-bin/hgTables? AluAluCNVpredictor: http://alualucnvpredictor.research.bcm.edu:3838/

gnomAD: https://gnomad.broadinstitute.org/

MCPS variant browser: https://rgc-mcps.regeneron.com/home

All of Us: https://databrowser.researchallofus.org/variants/

VizCNV: https://github.com/BCM-Lupskilab/VizCNV

## Acknowledgments

We thank the families, colleagues and collaborators for their participation in this study. We utilized ChatGPT to correct grammatical errors and to enhance the clarity of the manuscript.

## Availability of data and materials

The data generated or analyzed during this study are included in this published article. The released version (v4.2.3) of VizCNV and the processed 1000 Genome data can be accessed through https://doi.org/10.6084/m9.figshare.25869523[34]

## Authors’ contributions

Conceptualization: H.D., M.L., J.R.L and C.C; Data curation: S.N.J., S.V.B., D.M.M., M.L., C.M.G., L.G., M.G.M, M.C.P., I.K.C.; Formal Analysis: H.D., M.L., L.G.; Funding acquisition: J.R.L., C.C., R.A.G., J.E.P; Visualization: H.D., M.L.; Methodology: H.D., M.L., C.C. J.R.L.; Resources: M.C.P., I.K.C., J.R.L., J.E.P., R.A.G.; Supervision: C.C., J.R.L.; Writing – original draft: H.D.; Writing – review & editing: H.D., M.L., J.R.L., C.C. All authors reviewed the manuscript.

## Ethics approval and consent to participate

This study was performed with approval by the Institutional Review Board (IRB) at Baylor College of Medicine under research protocols H-29697 and H-42680.

## Consent for publication

Written consent was obtained to publish clinical information of the patients reported herein.

## List of abbreviations

aCGH: array Comparative Genomic Hybridization
AAMR: *Alu*-*Alu* Mediated Rearrangement
BAF: B-allele Frequency
CNV: Copy Number Variation
ES: Exome Sequencing
FDR: False Discovery Rate
GATK: Genome Analysis Toolkit
GS: Genome Sequencing
GREGoR: Genomics Research to Elucidate the Genetics of Rare Diseases
IGV: Integrative Genomics Viewer
PIDD: Primary Immunodeficiency Disease
ROH: Runs of Homozygosity
SD: Segmental Duplication
SV: Structural Variation
SNP: Single Nucleotide Polymorphism

---

**Additional file 1: Supplementary figure S1-S8.**

**Additional file 2: Table S1-S5.**

