## Additional file 1 for "VizCNV: An integrated platform for concurrent phased BAF and CNV analysis with trio genome sequencing data"

S1

a

b

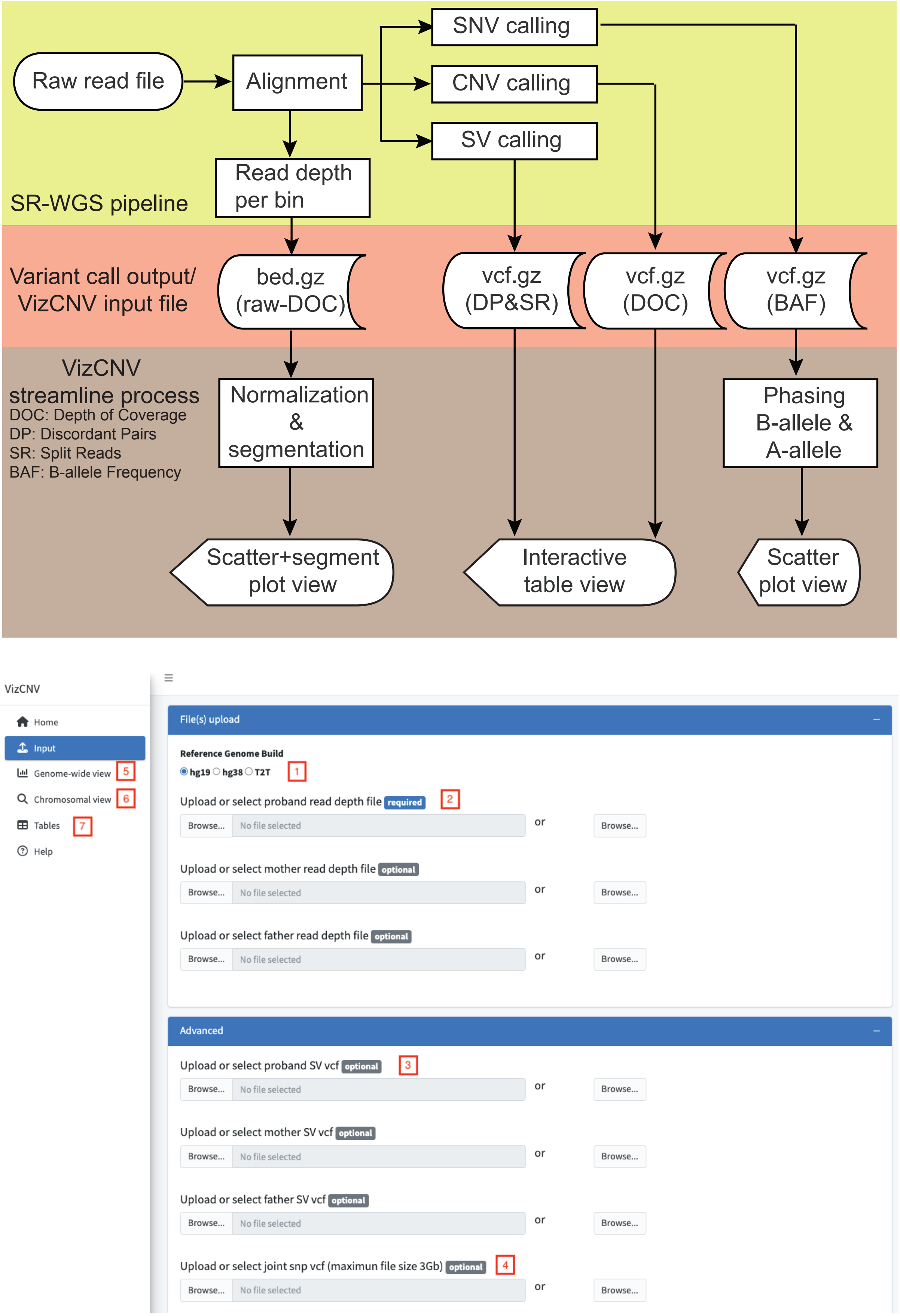

SNV calling
Raw read file
Alignment
CNV calling
SV calling
Read depth per bin
SR-WGS pipeline
Variant call output/ VizCNV input file
bed.gz (raw-DOC)
vcf.gz (DP&SR)
vcf.gz (DOC)
vcf.gz (BAF)
VizCNV streamline process
DOC: Depth of Coverage
DP: Discordant Pairs
SR: Split Reads
BAF: B-allele Frequency
Normalization & segmentation
Phasing B-allele & A-allele
Scatter+segment plot view
Interactive table view
Scatter plot view
VizCNV
Home
Input
Genome-wide view
5
Chromosomal view
6
Tables
7
Help
File(s) upload
Reference Genome Build
hg19 hg38 T2T
1
Upload or select proband read depth file required
2
Browse... No file selected
or
Browse...
Upload or select mother read depth file optional
Upload or select father read depth file optional
Advanced
Upload or select proband SV vcf optional
3
Upload or select mother SV vcf optional
Upload or select father SV vcf optional
Upload or select joint snp vcf (maximun file size 3Gb) optional
4

**Figure S1. The framework of VizCNV. a)** VizCNV workflow demonstrating the common output files from a standard short read genome sequencing pipeline incorporated into the VizCNV framework. **b)** User interface of the VizCNV. 1) Build-in reference. 2) input for read depth file. 3) input of proband structure variant call file. 4) joint-called genotype file. 5) genome-wide view for aneuploidy or sub-chromosomal events. 6) chromosome view for CNV and BAF inspection. 7) table view with annotate CNV information.

S2

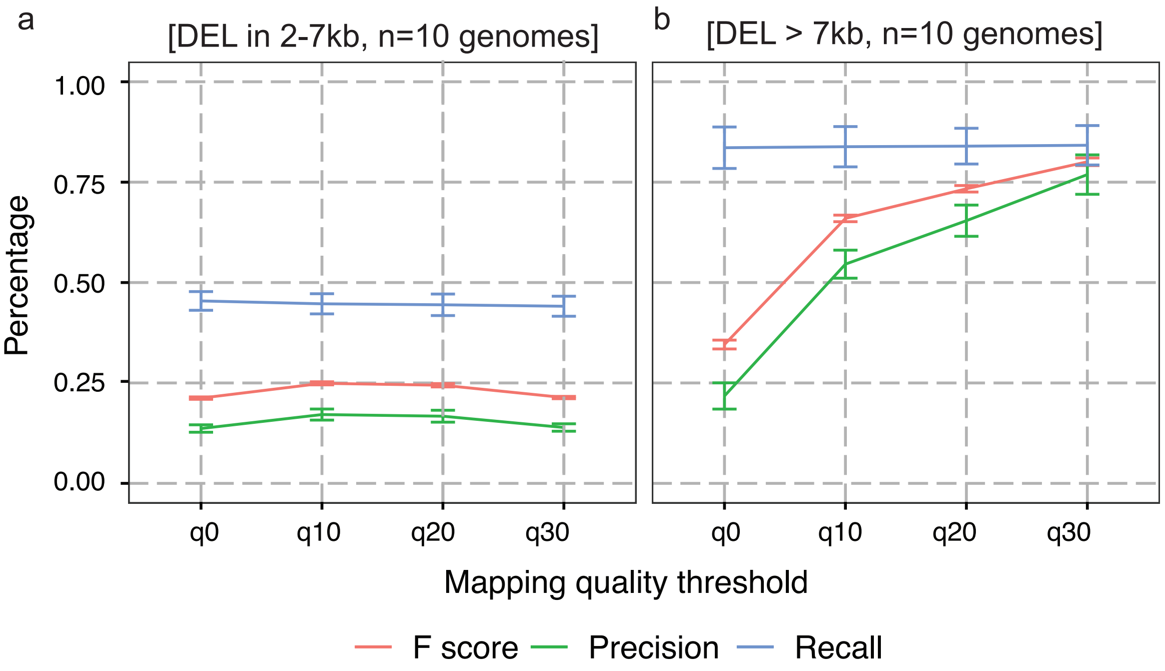
a
[DEL in 2-7kb, n=10 genomes]
b
[DEL > 7kb, n=10 genomes]
Percentage
1.00
0.75
0.50
0.25
0.00
q0
q10
q20
q30
Mapping quality threshold
F score
Precision
Recall

**Figure S2. Performance Metrics of Deletion Detection at Various Mapping Quality Thresholds across 10 1k genomes. a)** Deletions ranging from 2-7 kb. Metrics include F score (red), precision (green), and recall (blue), demonstrating the stability of the F score and precision across all thresholds, with a modest increase in recall from q0 to q30. **b)** Deletions larger than 7 kb. The F score, precision, and recall all show an upward trend with increasing mapping quality thresholds, indicating improved detection performance at higher thresholds.

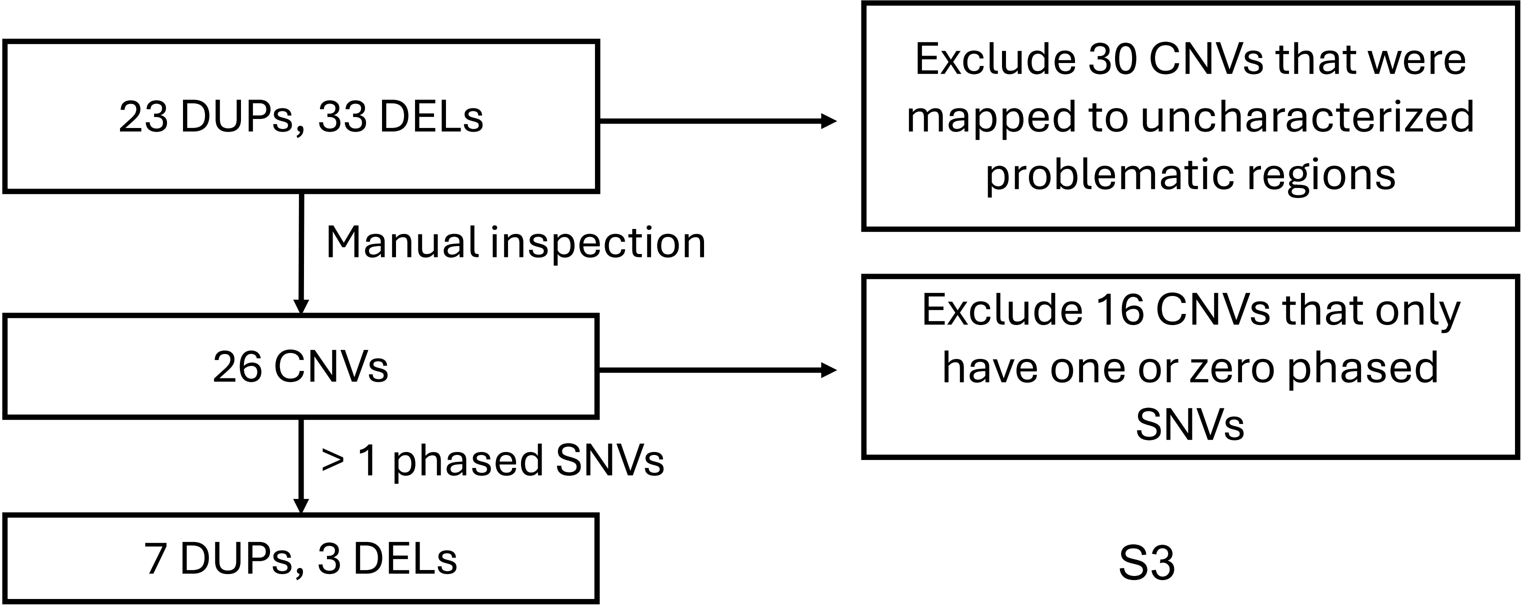
23 DUPs, 33 DELs
Exclude 30 CNVs that were mapped to uncharacterized problematic regions
Manual inspection
26 CNVs
Exclude 16 CNVs that only have one or zero phased SNVs
> 1 phased SNVs
7 DUPs, 3 DELs

S3

**Figure S3. Schematic Representation of Joint BAF and Read Depth Analysis in the HG00405 Case.**

S4

HG00405, HGSV_211297, chr16:78673486-79733774

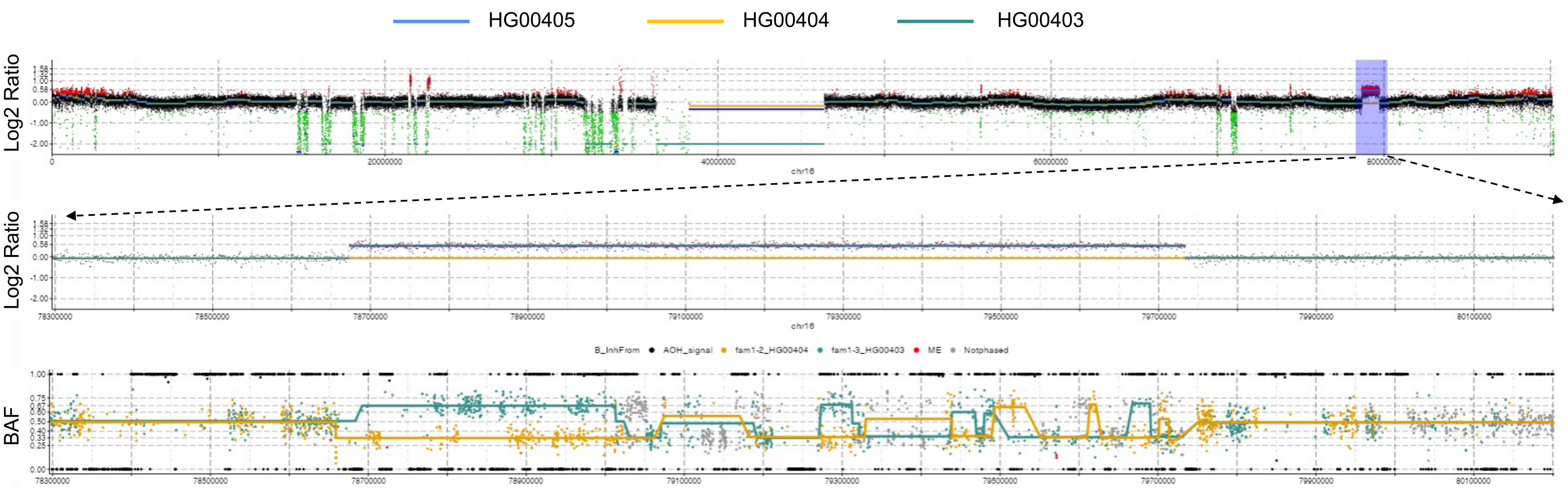

**Figure S4. Paternal Inheritance of Duplication with Potential Recombination.** The top panel presents a whole chromosome $\log_2$ ratio plot for the child (HG00405), father (HG00403), and mother (HG00404), with a blue rectangle highlighting a ~1 Mb duplication identified in both the child and father. The middle and bottom panels offer a detailed view of the highlighted region, focusing on $\log_2$ ratio and B-allele frequency (BAF), respectively. Colors indicate the origin of the variant: teal for paternal and orange for maternal.

S5

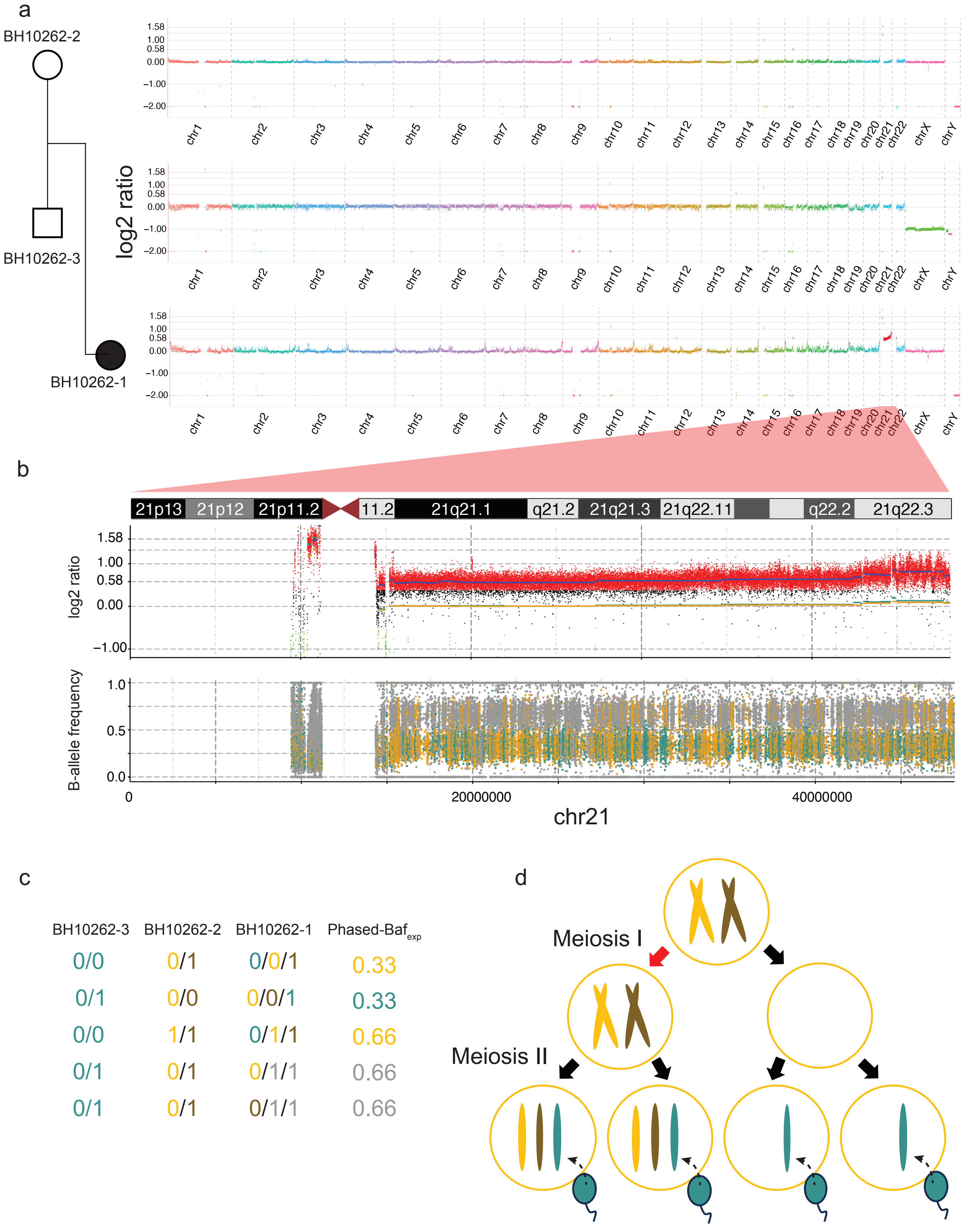
a
BH10262-2
BH10262-3
BH10262-1
log2 ratio
1.58
1.00
0.58
0.00
−1.00
−2.00
chr1 chr2 chr3 chr4 chr5 chr6 chr7 chr8 chr9 chr10 chr11 chr12 chr13 chr14 chr15 chr16 chr17 chr18 chr19 chr20 chr21 chr22 chrX chrY
b
21p13 21p12 21p11.2 11.2 21q21.1 q21.2 21q21.3 21q22.11 q22.2 21q22.3
log2 ratio
B-allele frequency
1.0
0.5
0.0
0
20000000
40000000
chr21
c
BH10262-3 BH10262-2 BH10262-1 Phased-Baf_exp
0/0 0/1 0/0/1 0.33
0/1 0/0 0/0/1 0.33
0/0 1/1 0/1/1 0.66
0/1 0/1 0/1/1 0.66
0/1 0/1 0/1/1 0.66
d
Meiosis I
Meiosis II

**Figure S5. Genomic analysis of a PIDD family with trisomy 21. a)** Genome-wide $log_2$ ratio plot with pedigree showing on the left. **b)** Detailed $log_2$ ratio and B allele frequency plots for chromosome 21 of the proband, BH10262-1. The top panel shows the $log_2$ ratio with regions of interest highlighted, and the bottom panel shows phased B allele frequency- color coding indicates allele origins: yellow for from maternal origin (BH10262-2), blue for paternal (BH10262-3), and grey for indeterminate. **c)** Table show the schema of the phasing. **d)** Schematic representation of meiotic divisions I and II, illustrating chromosomal behavior and potential segregation patterns that could lead to the observed genetic variations in the family. Chromosome coloring corresponds to the B allele origins in **b and c.**

S6

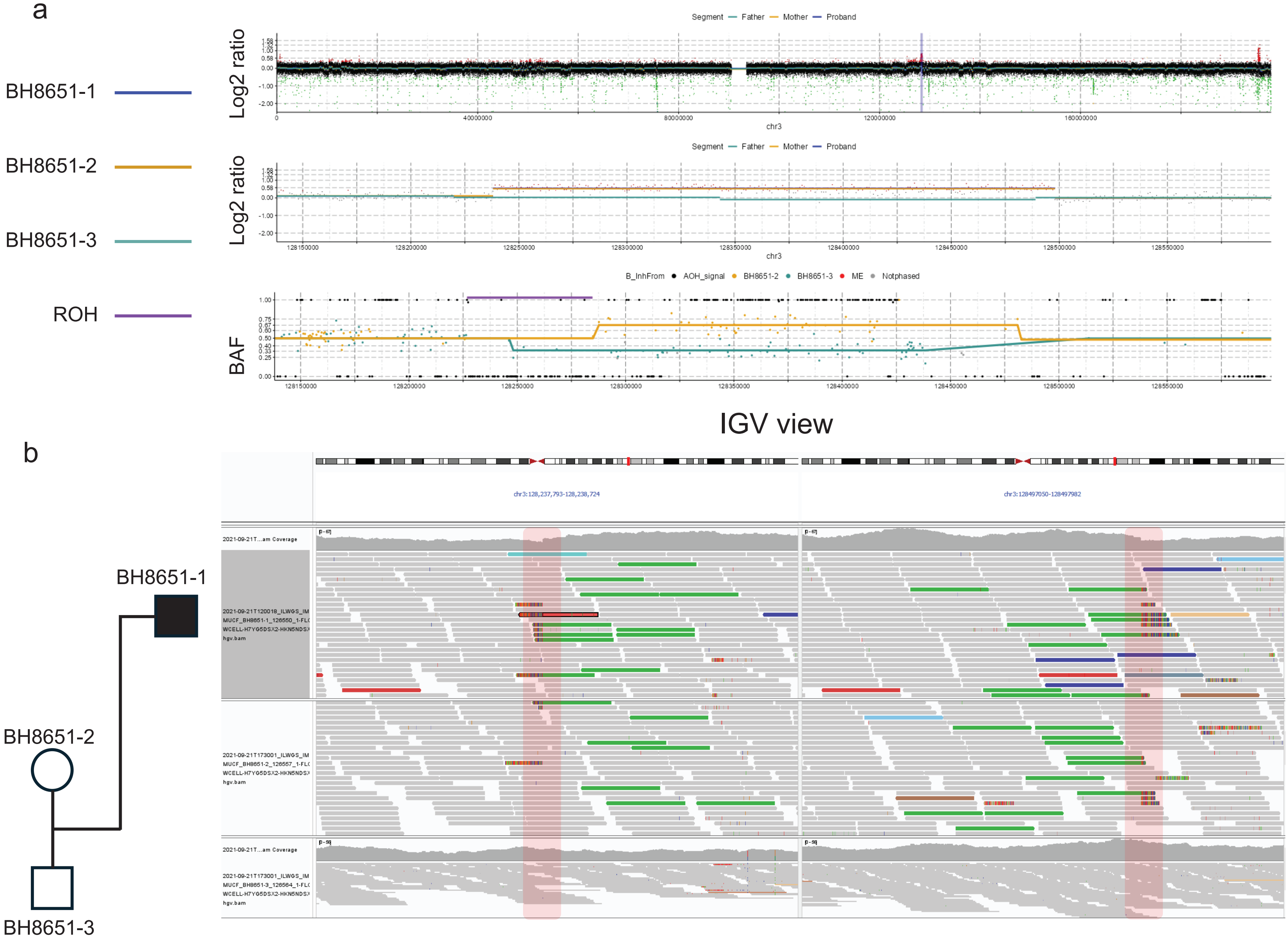

a
VizCNV view
BH8651-1
BH8651-2
BH8651-3
ROH
Segment — Father — Mother — Proband
Log2 ratio
chr3
B_InhFrom ● AOH_signal ● BH8651-2 ● BH8651-3 ● ME ● Notphased
BAF
b
IGV view
BH8651-1
BH8651-2
BH8651-3
chr3:128,237,793-128,238,724
chr3:128497050-128497982

**Figure S6. Representative Duplication with Consistent BAF and $log_2$ Analysis. a)** The top panel displays the $log_2$ ratio of chromosome 3. The middle panel provides a zoomed-in view of the duplication region, while the bottom panel shows the BAF, with lines representing segmented BAF values. Colors represent segments from family members: blue for the proband, teal for the father, and yellow for the mother. Purple segments denote region with ROH call. **b**) The left side shows the pedigree, and the right side presents the IGV view of the duplication. Highlighted regions point to changes in read depth and several discordant read pairs suggesting a tandem duplication in both the proband and the mother. This case presents a maternally inherited duplication with identical haplotypes.

S7

VizCNV view

a

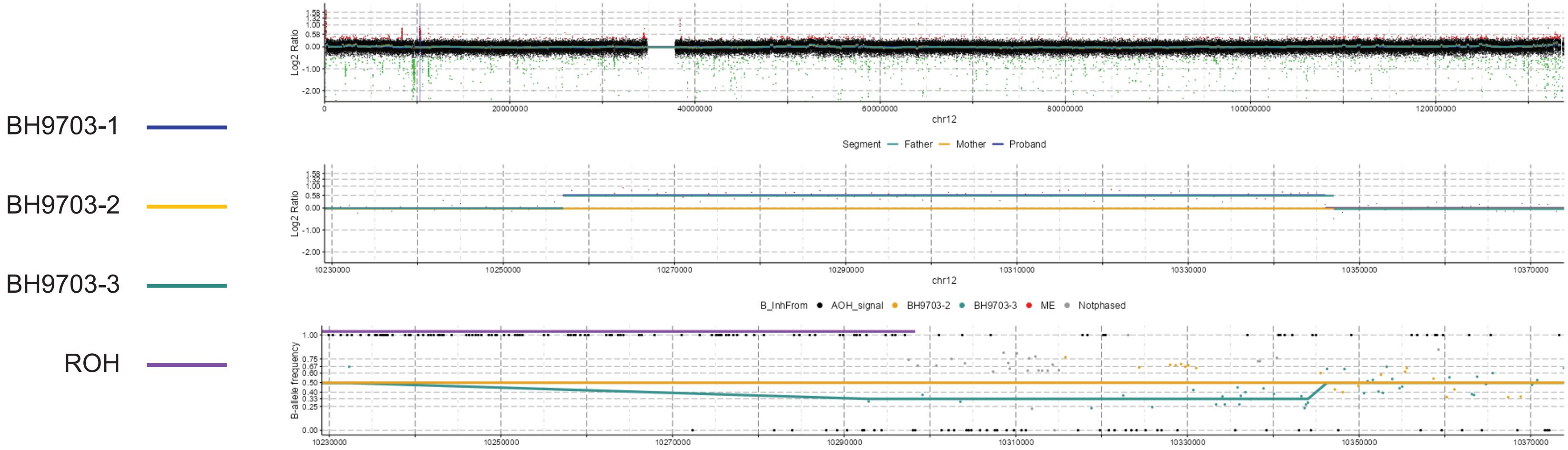


IGV view

b

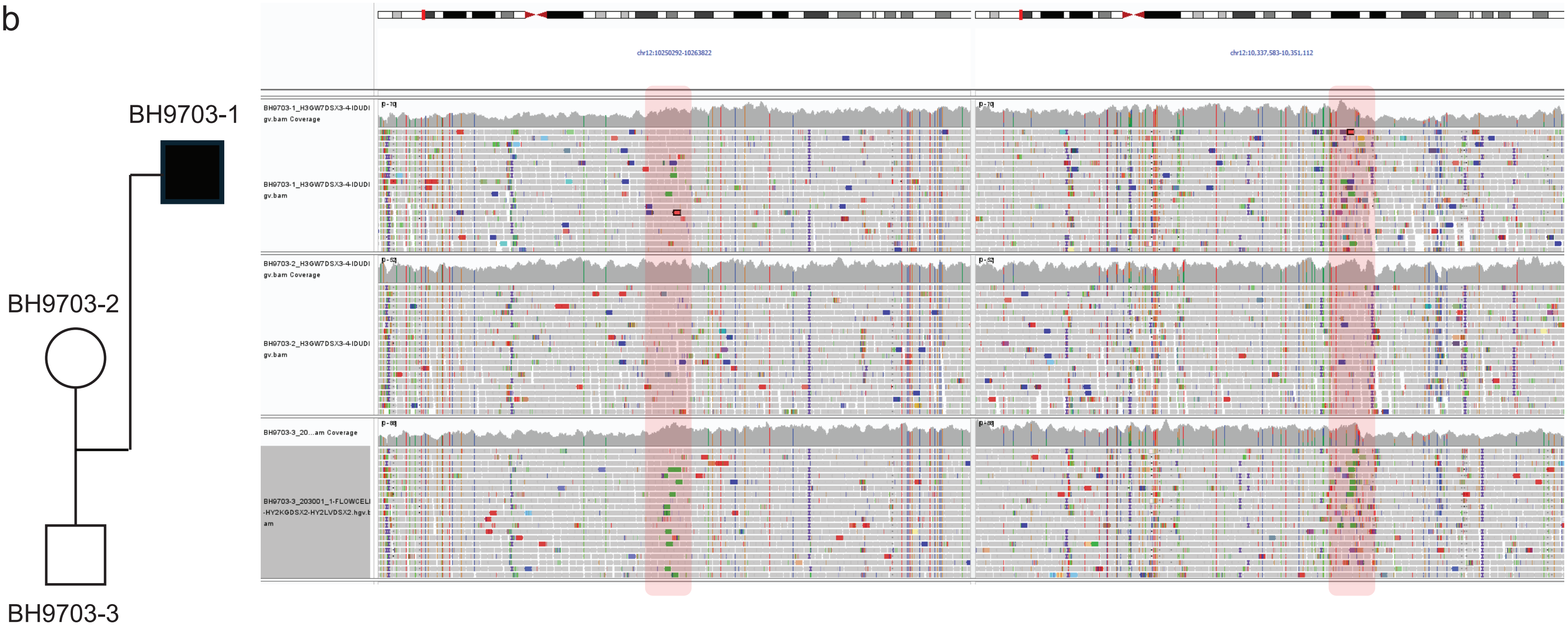

**Figure S7. Paternally Inherited Duplication with Two Different Haplotypes. a**) The top panel displays the $\log_2$ ratio of chromosome 12. The middle panel provides a zoomed-in view of the duplication region, and the bottom panel shows the BAF, with lines representing segmented BAF values. Colors denote individuals from the family: blue for the proband, teal for the father, and yellow for the mother. **b**) The left side shows the pedigree, and the right side presents the IGV view of the duplication. Highlighted regions indicate changes in read depth and several discordant read pairs, suggesting a tandem duplication in both the proband and the father.

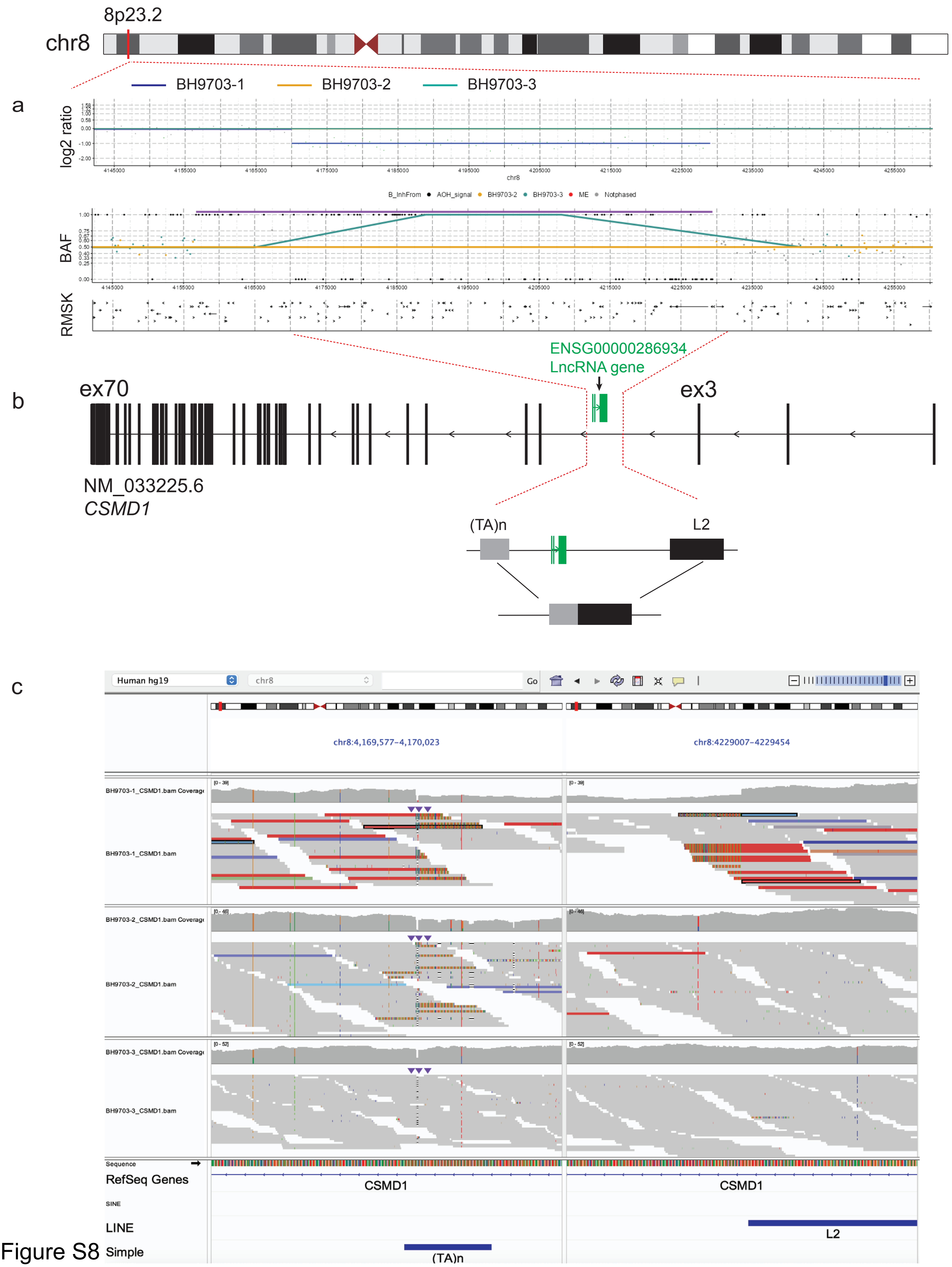
8p23.2
chr8
BH9703-1
BH9703-2
BH9703-3
a
log2 ratio
BAF
RMSK
ENSG00000286934
LncRNA gene
ex70
ex3
b
NM_033225.6
CSMD1
(TA)n
L2
c
Human hg19
chr8
Go
chr8:4,169,577-4,170,023
chr8:4229007-4229454
BH9703-1_CSMD1.bam Coverage
BH9703-1_CSMD1.bam
BH9703-2_CSMD1.bam Coverage
BH9703-2_CSMD1.bam
BH9703-3_CSMD1.bam Coverage
BH9703-3_CSMD1.bam
Sequence
RefSeq Genes
CSMD1
SINE
LINE
Simple
L2
(TA)n

Figure S8

**Figure S8.** ***De Novo*** **Deletion of LncRNA in Proband BH9703-1. a)** The upper panel illustrates the ideogram of chromosome 8. The lower panel presents VizCNV output, displaying $log_2$ ratio and B-allele frequency plots that identify a *de novo* deletion impacting *CSMD1*. **b)** The upper panel displays the canonical *CSMD1* transcript along with a smaller noncoding gene (highlighted in green), which is antisense to *CSMD1*. The deletion, within intron 3 of *CSMD1*, does not disrupt any exons but completely remove the entire long non-coding RNA gene. The diagram also provides a detailed view of the deletion's genomic structure. Noting the simple repeat and LINE element near the breakpoint. **c)** IGV visualization of the deletion's breakpoint junction of SR-WGS.
